# Syllable repetition reveals links between distant phrases in birdsong

**DOI:** 10.64898/2026.05.12.724669

**Authors:** Priya Binwal, Lena Veit

## Abstract

Repeated execution of individual behavioural units is a common feature of many learned motor behaviours such as dance, music, and birdsong. Little is known about the neuronal control of such learned motor sequences, and specifically, how the number of variable repetitions is determined. The songs of Bengalese finches (*Lonchura striata domestica*) consist of individual syllables which can repeat a variable number of times (repeat number) to form a repeat phrase. Like vocal sequences in other animals, Bengalese finch song syntax is typically modelled as a Markov chain, where the choice to repeat the same syllable type or switch to a different one is made stochastically after each syllable, before the next syllable is produced. Here, we report that repeat numbers of adjacent and distant repeat phrases in the song can be correlated across specific pairs of phrases. These hidden links between distinct phrases challenge existing models of song syntax where the number of repetitions is independently determined for each syllable type. Instead, they suggest an organisation where a joint factor can control multiple nonadjacent phrases in a song. Repeat phrases in Bengalese finches may therefore be particularly suited to study the neuronal mechanisms underlying long-range dependencies in complex vocal sequences.

## Introduction

Multiple repetitions of the same element are common in complex behaviours such as music, dance, and other vocal and non-vocal communication sequences in humans and other animals [1–9]. For example, many birdsongs contain repeat phrases or trills, where the same syllable type is repeated a variable number of times [10–19]. Like speech, birdsong is a learned vocal behaviour controlled by analogous forebrain circuits, and relies closely on sensory feedback [20–22]. Therefore, birdsong is a useful model to study the neural mechanisms involved in the production of variably repeating vocal units.

The songs of male Bengalese finches (*Lonchura striata domestica*) contain both repeating and non-repeating syllable types which occur in variable order. Syllable production involves the activation of premotor networks in the nucleus HVC (proper name), and probabilistic transitions between syllables are typically modelled as a Markov process relying only on syllable type and recent song history [23–26]. After the production of each syllable, the next syllable is thought to be chosen through a stochastic winner-take-all mechanism between premotor networks encoding each syllable, that plays out during the gap before the next syllable [23,24,27,28]. To explain observed repeat count distributions of repeat phrases, which are inconsistent with a pure Markov mechanism [24,28], this model of song syntax has been extended to incorporate adapting self-transitions. The adaptation of self-transition probabilities is driven by adapting auditory feedback with declines based on an adaptation parameter that is specific to each syllable type [28]. Such a model fits existing repeat count distributions well. Thus, the presumed mechanism of repeat production posits that the number of times a syllable repeats within a phrase is specific to that syllable type, with repetitions of different syllable types being controlled by independent parameters.

We here turn our attention to the question: which factors determine the number of times a given syllable is repeated in Bengalese finch songs? We show that the number of times syllables repeat within repeat phrases is influenced by the identity of preceding or following syllables, consistent with Markov models. However, we also find local and non-local dependencies between pairs of repeating syllables which are not captured by current syntax models, suggesting the presence of higher order structure in Bengalese finch song.

## Materials and Methods

### Subjects & Sound recording

Songs were recorded from 6 adult male Bengalese finches (*Lonchura striata domestica*). Four birds were recorded at the University of California, San Francisco, and two birds were recorded at the University of Tübingen, Germany. All birds were adults (>120 days post hatch, dph) at the time of recording (mean age ∼286 dph, range 132-691 dph for four birds for which the exact age was known). Birds were placed in sound-attenuating boxes and recorded alone for multiple days (mean 3.5, range 2-6), and a 14:10 hour light:dark period was maintained. Sound was recorded using custom LabView software or Moove [29,30]. On average, we included ∼622 song bouts (range 300-985) from each individual in our analyses. All experiments were performed in accordance with animal protocols approved by the University of California, San Francisco Institutional Animal Care and Use Committee (IACUC) or the national authority, the Regierungspräsidium Tübingen, Germany.

### Song annotation

A song bout was defined as a continuous period of song separated by 2s of silence as in [31,32]. Song bouts were semi-automatically annotated as described previously [29,33]. We hand-checked annotated labels to correct any errors. Consecutive occurrences of the same syllable were defined as a repeat phrase [30,31]. The labelled sequences were simplified by collapsing repeat phrases of a given type into a single state, merging the introductory notes and inter-notes into a single state ‘I’, and merging stereotyped sequences or chunks into a single state [31,32]. Thus, a single state in our simplified sequence could represent an introductory or inter-note, a repeat phrase, single syllables, or a chunk. We obtained transition probabilities between these states and represented them in transition diagrams as in **Figure 2E**.

### Repeat distributions in different sequential contexts

We used a k-sample Anderson Darling test to test whether the repeat distribution of a phrase was significantly affected by at least one context (preceding or upcoming phrase in the bout). We excluded contexts that were rarer than 5% from our analyses. Distributions for different preceding and upcoming contexts were compared separately.

### Correlations between repeat phrases

We analysed relationships between repeat numbers of repeat phrase pairs in the same song bout using Spearman’s rank correlation coefficients, followed by the Benjamini-Hochberg correction for multiple comparisons. For a given pair of repeat phrases A and B, this was implemented in two ways: 1. By selecting adjacent occurrences of A and B in all song bouts and calculating the correlation coefficient between their repeat numbers. A threshold of a minimum 100 adjacent occurrences was used to exclude repeat pairs that rarely occurred together. 2. By selecting all occurrences of A and the next occurrence of B (irrespective of their distance in the song bout) and calculating the correlation coefficient between their repeat numbers. If there were multiple occurrences of B that followed A in the song bout, only the first following B was used. In both cases, all analyses were restricted to phrases occurring within the same song bout.

To analyse whether distance between repeat phrases affects the observed correlations, we defined distance in units of phrases, as depicted in **Figure 2**; therefore, directly adjacent phrases will have a distance of 1. For a given pair of repeat phrases A and B, we calculated the correlation between their repeat numbers at all possible distances up to 21. A threshold of a minimum of 100 occurrences at a given distance was used to exclude rarely occurring distances. All correlations were tested directionally, i.e. the correlation between A and B is not the same as the correlation between B and A, as the latter would be calculated on instances where B occurred before A in the bout.

We relied on a bootstrapping approach to ensure that the observed correlations were not a byproduct of constraints imposed by the repeat distributions. We first converted the song sequence into a compressed format where letters represent phrases in the song with a number following them to represent their repeat numbers. For example, a song sequence like aaabbccccd was represented as A_3_B_2_C_4_D_1_. Following this, we created a simulated dataset by keeping the phrase order in the bout intact and sampling the repeat number of each phrase from its own repeat distribution. For example, A_3_B_2_C_4_D_1_ was used to create A_na_B_nb_C_nc_D_nd_ where n_a_, n_b_, n_c_, and n_d_ were randomly sampled from the repeat distributions of A, B, C and D respectively. In this way, each song bout in our original dataset was used to simulate one song bout for the simulated dataset.

We calculated correlations between pairs of repeat phrases in the simulated data exactly as described for the real data. This process was repeated for 100 simulated datasets resulting in 100 values of expected correlation coefficients for each repeat phrase pair. The observed correlation between each pair was compared against the distribution of expected correlations for that pair with a one-tailed z-test to test the hypothesis that the observed correlation coefficient was greater than what we would expect by chance given the repeat distributions of repeat phrases (**Figure S1**). Only correlations that were significant after the first p-value correction were tested this way. All adjacent pairs analysed (13 out of 13; 100%) remained significant after this analysis while 40 out of 47 significant correlations between all phrases (including non-adjacent ones) remained significant (85.1%).

### Effect of distance on correlations between repeat numbers

We analysed the effect of distance on correlations between repeat numbers of phrase pairs, including self-pairs, using a linear mixed effect model (LMM) which included distance as a fixed effect and syllable pair nested within bird identity as a random effect.

### Analysis of repeat number elasticities

If *L* represents song bout length (number of syllables in the bout) and *n_i_* represents the repeat number of a repeat phrase, then we can write the following regression equation:

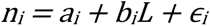

where, *a_i_* is the intercept, *b_i_* is the regression coefficient and *ɛ_i_* is the residual. The regression coefficient *b_i_* represents how the repeat number of a given phrase changes in proportion to changes in bout length. To make *b_i_* comparable across repeat phrases and birds, we normalised it by dividing it by the value of the regression coefficient that is expected if proportional scaling holds [34] as follows:

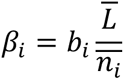

*β*_i_ is termed as the elasticity of a repeat phrase and it measures how much the repeat number of a repeat phrase changes in proportion to changes in bout length. More specifically, if *β*_i_ = 1, the repeat number of the repeat phrase changes in proportion to bout length. *β*_i_ > 1 and *β*_i_ < 1 would imply the repeat number of the repeat phrase scales more or less than what would be expected by proportional scaling respectively, while *β*_i_ = 0 would mean that the repeat number of the repeat phrase is invariant to changes in bout length. This approach was adapted from a study in zebra finches by Glaze and Troyer [34].

### Acoustic features

We used the audio feature extractor function from MATLAB’s Audio Toolbox (Version 24.2), to extract 31 acoustic features for each occurrence of each syllable [35, 36]. For each bird, we performed a PCA for dimensionality reduction and included as many PCs as were needed to explain at least 90% of the variance in further analyses (ranged from first 7-13 PCs across birds). To quantify acoustic distances between syllables, we calculated Euclidean distances between centroids of already labelled syllable clusters in PC space. The acoustic distances we obtained were z-scored to make them comparable across birds.

To compare acoustic distances between positively correlated, negatively correlated, and uncorrelated repeat phrases, an LMM with bird identity as a random effect followed by pair-wise post-hoc tests using estimated marginal means [37] with a Tukey-correction for p-values was used.

When analysing correlations between adjacent occurrences of repeat phrases such as AB, the calculation of acoustic distance between A and B was based on only those instances of A and B that occurred adjacently in the order AB. This was done to ensure that our measure of acoustic difference was not affected by context-dependent differences in syllable acoustics [38]. However, when analysing correlations between A and the next occurrence of B (i.e. including non-adjacent occurrences), we relied on acoustic distances calculated based on all occurrences of A and B.

### Position in song bout

We studied the effect of position in song bout on repeat numbers by testing: 1. Whether later occurrences of the same repeat phrase in the bout have smaller repeat numbers in comparison to earlier occurrences. 2. Whether repeat phrases have smaller repeat numbers later in the bout in comparison to the beginning of the bout. Note that the second possibility doesn’t necessarily imply the first— for example, it is possible for the same repeat phrase to occur twice near the beginning of the song bout and differ in repeat numbers. We used linear regression to analyse whether the repeat numbers change with occurrence number (i.e. whether the phrase occurred for the first time, second time etc. in the song bout) or relative position in the bout (start position of the repeat phrase divided by bout length). Thus, two separate linear regressions were carried out to study the effect of position in bout on each repeat phrase. The slope of the regression line was standardised by dividing by the standard deviation of the corresponding repeat distribution.

### Time of day

To analyse the effect of the time of day on repeat distributions, we fit a linear regression to repeat numbers of each repeat phrase in our dataset against time of day (hour of recording between 6am-10pm). The slopes of the regression line were standardised by dividing by the standard deviation of the corresponding repeat distribution.

### Random Forest models

We used random forest models to examine the combined effects of different factors that could influence repeat numbers. A separate random forest regression model was trained to predict the repeat numbers of each repeat phrase based on the identities and repeat numbers of four preceding and upcoming phrases in the song. In addition, bout length, occurrence number, relative position of the repeat phrase, and time of day were also included as features. All categorical features (identities of previous and upcoming phrases) were one-hot encoded, and the data were split into training and validation sets using five-fold cross-validation. A grid-search was implemented to tune the hyperparameters (number of trees, maximum depth of a tree, and maximum number of features to be considered at each split) of each model. We implemented a threshold of a minimum sample size of 1000. An OOB (out of the bag score) was calculated for each model and models for which the difference between the OOB score and the cross-validated R^2^ was more than 0.1 were excluded to exclude overfitted and underfitted models. Of the 23 repeat phrases in our dataset, 16 phrases from 5 birds survived these filtering criteria. Feature importances were assessed using SHAP values [39]. Since categorical features were one-hot encoded, this resulted in separate SHAP values by phrase type for each such feature (for e.g., preceding phrase A, preceding phrase B etc.). Taking advantage of the additive property of SHAP values, we aggregated these values for each categorical feature. For example, if the repeat phrase being modelled has two possible preceding phrase types A and B, then the SHAP value of the feature ‘preceding phrase’ was simply calculated as the sum of SHAP values of features ‘preceding_A’ and ‘preceding_B’. SHAP values are in units of the response variable, i.e. in units of repeat numbers in this case. We divided the SHAP values of each phrase-specific model by the standard deviation of the repeat number of the modelled repeat phrase to obtain SHAP importances in units of standard deviation.

## Results

### Repeat distributions are affected by sequence context

The song of Bengalese finches (*Lonchura striata domestica*) consists of a limited number of acoustically discrete syllable types, which can be variably repeated in repeat phrases (**Figure 1A**; [40]). It is typically modelled as a first-order Markov sequence, with stochastic decisions for the next syllable playing out at the end of each syllable, influenced by its identity and recent history [23–28]. We found that distributions of repeat numbers (how often syllables were produced in each phrase), depended on syllable identity and could additionally depend on the sequence context of preceding or upcoming syllable types (**Figures 1C-F**). For all tested phrases occurring in multiple contexts (n=15 phrases from 6 birds), the upcoming context significantly affected repeat distributions for 14/15 phrases (93.8%; p < 0.05, Anderson Darling test; **Figure 1G**), the preceding context significantly affected repeat distributions in all tested phrases (n=12 phrases from 6 birds; **Figure 1G**; p < 0.05, Anderson Darling test). Thus, the identities of both preceding and upcoming phrases can influence repeat number distributions, consistent with partially observable Markov models, where multiple states can produce the same syllable type with different repeat distributions [24,25,41].

**Figure 1:**
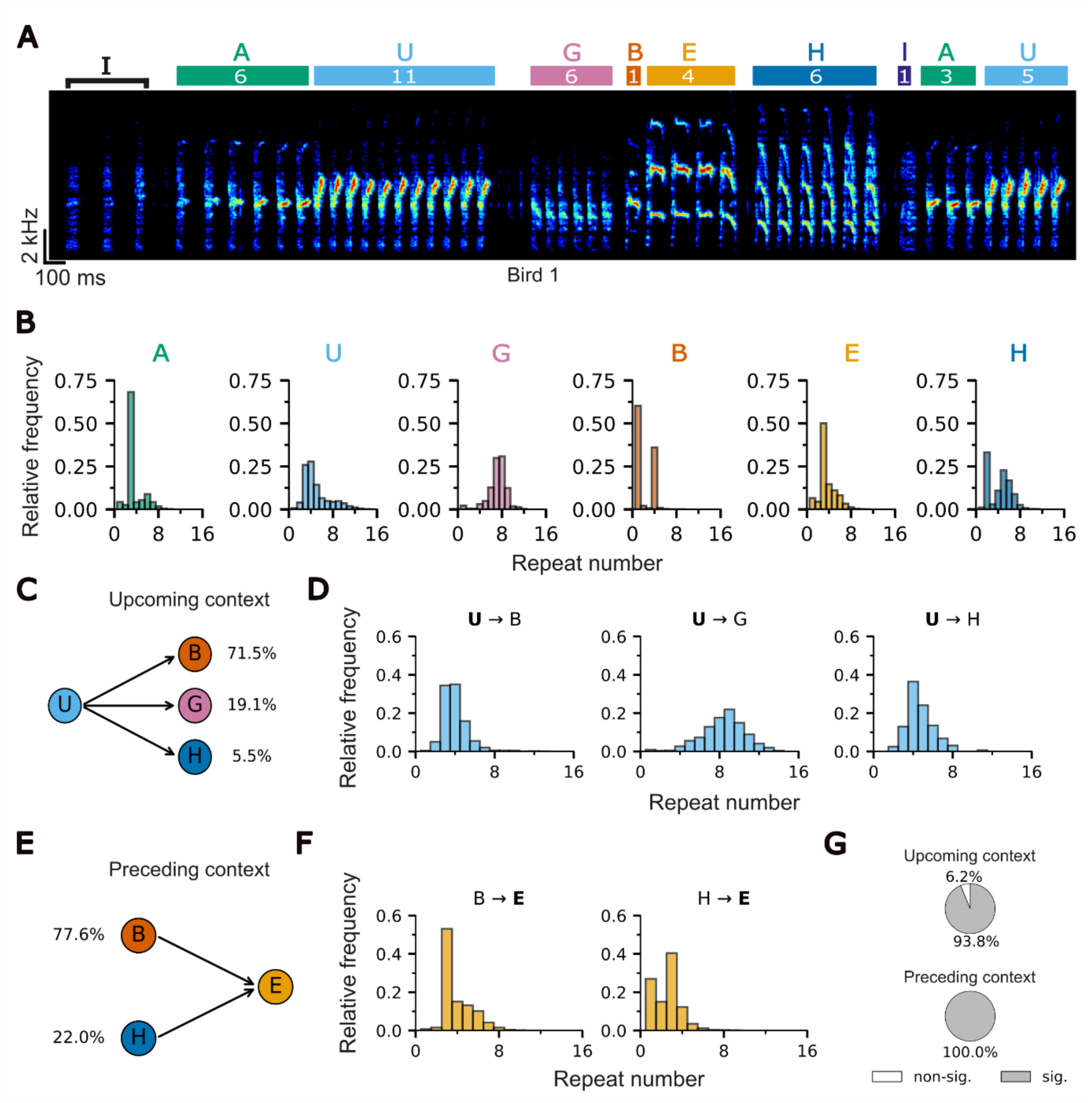
Repeat number distributions depend on sequence context. (A) Spectrogram of a Bengalese finch song where letters indicate phrase types. ‘I’ denotes the introductory state. Uppercase and lowercase letters denote repeat phrases and non-repeating syllables respectively. Coloured rectangles indicate the extent of each phrase with numbers inside denoting repeat numbers. (B) Repeat distributions of all repeat phrases shown in panel A. (C) Three possible upcoming contexts for the example phrase U shown in panels A, B. Percentages indicate probabilities of transitioning from U to the next phrase. (D) Repeat distributions of U in three different upcoming contexts. (E) Two possible preceding contexts for the example phrase E shown in panels A, B. Percentages indicate probabilities of E being preceded by these phrases. (F) Repeat distributions of E in two different preceding contexts. (G) Pie charts indicate the proportion of phrases with significantly different repeat distributions (k-sample Anderson Darling test) for at least one upcoming (top) or preceding (bottom) context.

### Repeat numbers of specific pairs of phrases are correlated

We next investigated whether not just the identity, but also the repeat number of surrounding phrases can influence repeat number distributions. **Figures 2A, B** show a particularly striking example of neighbouring phrases occurring with similar repeat numbers across bouts (Spearman’s *ρ* =0.61). Across our dataset of six birds, we found significantly positive correlations in 28.6% of adjacent phrase pairs and significantly negative correlations in 17.9% of adjacent pairs (**Figure 2C**; n = 28 pairs from 6 birds). The average magnitude of adjacent correlations was similar for positively correlated and negatively correlated pairs (0.31 ± 0.09 and −0.35 ± 0.11, respectively).

**Figure 2:**
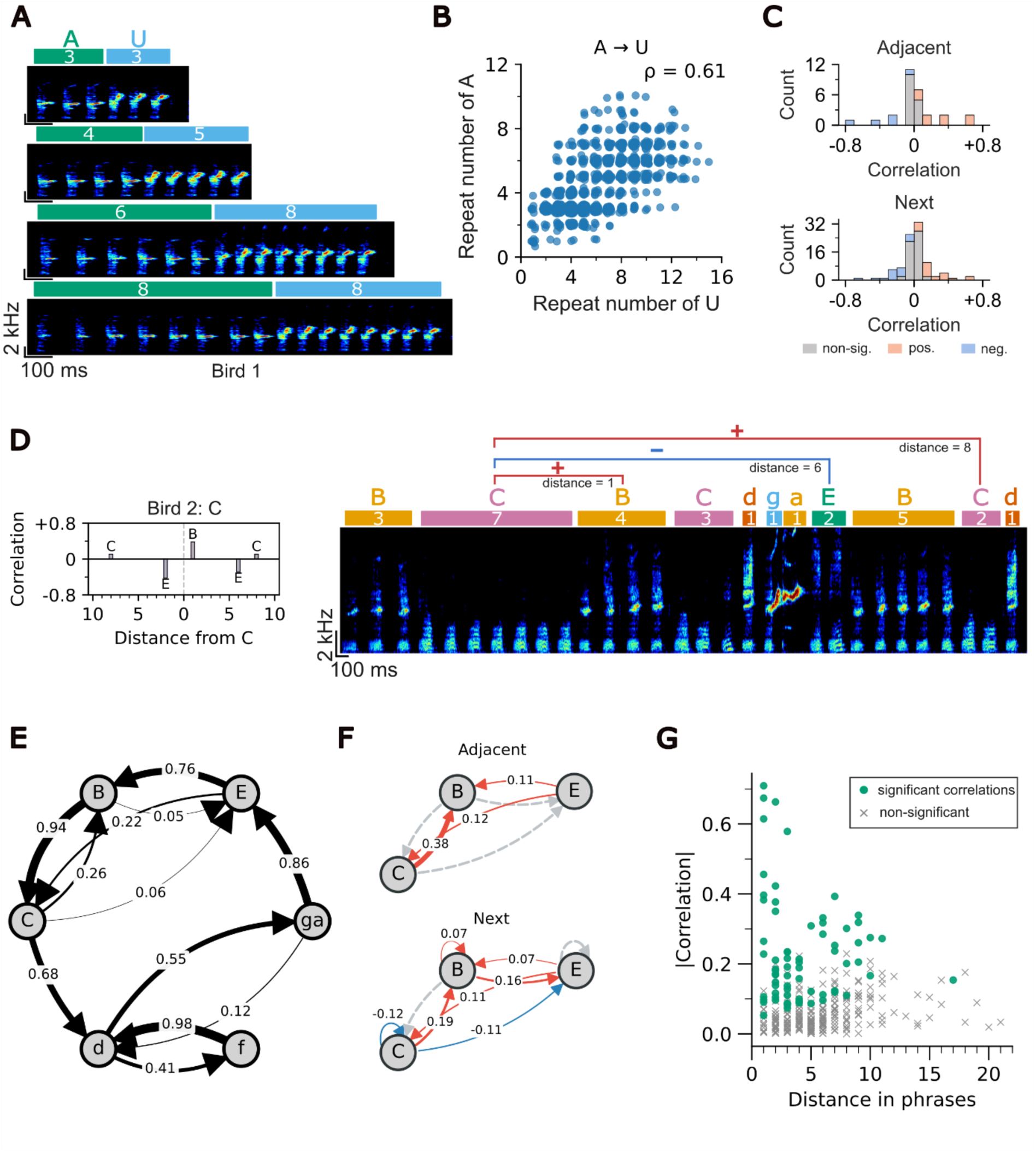
Repeat numbers of specific phrase pairs are correlated. (A) Spectrograms from an example bird depict short and long A phrases followed by short and long U phrases respectively. (B) Scatterplot depicts a positive correlation between repeat numbers of adjacent phrases A and U. (C) Histogram counts of positive, negative, and non-significant correlations for adjacent (top) and all phrase pairs (bottom). Distributions significantly deviated from normal (Shapiro-Wilk test for normality: W= 0.92, p<0.001). (D) Left: correlations between repeat numbers of different repeat phrase types with example phrase C at different distances. Right: Example spectrogram with correlations between C and subsequent phrases B, E, and C at different phrase distances. (E) Transition diagram for the example bird shown in D. Nodes indicate different states and numbers on edges indicate transition probabilities between states. Repeat phrases and non-repeating syllables are in uppercase and lowercase respectively. The chunk ‘ga’ is depicted as a single node. (F) Correlation diagrams depict links between adjacent and all pairs of repeat phrases analysed. Positively and negatively correlated phrase pairs are indicated by red and blue arrows respectively. Uncorrelated pairs are indicated by grey arrows. Self-correlations (circular arrows) are calculated between one phrase and a subsequent phrase of the same type. (G) Absolute value of correlations between repeat numbers of phrase pairs across different distances in the song bout (n=321 for 44 unique phrases occurring at different distances, range 1-7).

Do correlations only arise in directly adjacent phrases? We analysed correlations between repeat numbers of each phrase type and the next occurrence of another phrase type in the same bout, which may or may not be adjacent. In this larger dataset of 94 phrase pairs, we found similar proportions of correlated phrases (23.4% positive, 19.1% negative; **Figure 2C**).

**Figure 2D** shows an example song bout for bird 2, with links between phrases highlighted. As is typical for Bengalese finch songs, phrases can occur in variable order, thus the same pair can be observed at various distances (**Figure 2E**). This bird exhibited correlations between specific pairs of adjacent and non-adjacent phrases, while other pairs showed no correlations (**Figure 2F**). Across birds, the absolute value of the correlation coefficient was not significantly related to distance between phrase pairs (**Figure 2G**; p = 0.19 LMM; n = 44 unique pairs analysed from 6 birds, of which 14 pairs occurred at more than one distance), with a trend towards smaller magnitude correlations at higher distances.

Among all repeat phrase pairs that could be analysed in both directions (i.e. both the A→B and B→A correlation could be tested, n = 20), only 20% of correlations were symmetric (both directions positive or negative, 4/20 pairs), while 20% were anti-symmetric (positive correlation A→B and negative correlation B→A or vice versa, 4/20 pairs). The remaining pairs only exhibited unidirectional correlations. Therefore, specific pairs of phrases were correlated in specific contexts across the bout, exhibiting dependencies which are not captured by current sequence models.

### Repeat numbers are not strongly influenced by variations in song length

Correlations between repeat numbers of phrases in the same bout could be explained if there are global factors affecting the length of phrases [42]. For example, if all or some phrases grow longer or shorter in proportion to changes in overall bout length, this would lead to positive correlations between those phrases. We did not find such a relationship. Longer song bouts typically include additional phrases, rather than systematically longer phrases, with elasticity values ([34], see Methods) ranging from −0.25 to +0.11. This indicates that only a small proportion of variation in repeat numbers could be explained by scaling proportionally with bout length (**Figures 3A, B**). Additionally, we did not find a consistent pattern of linked phrases sharing the same elasticity sign for positive correlations or opposite elasticity signs for negative correlations, which could indicate bout length as a contributor to the observed correlations (**Figures S2A, B**).

**Figure 3:**
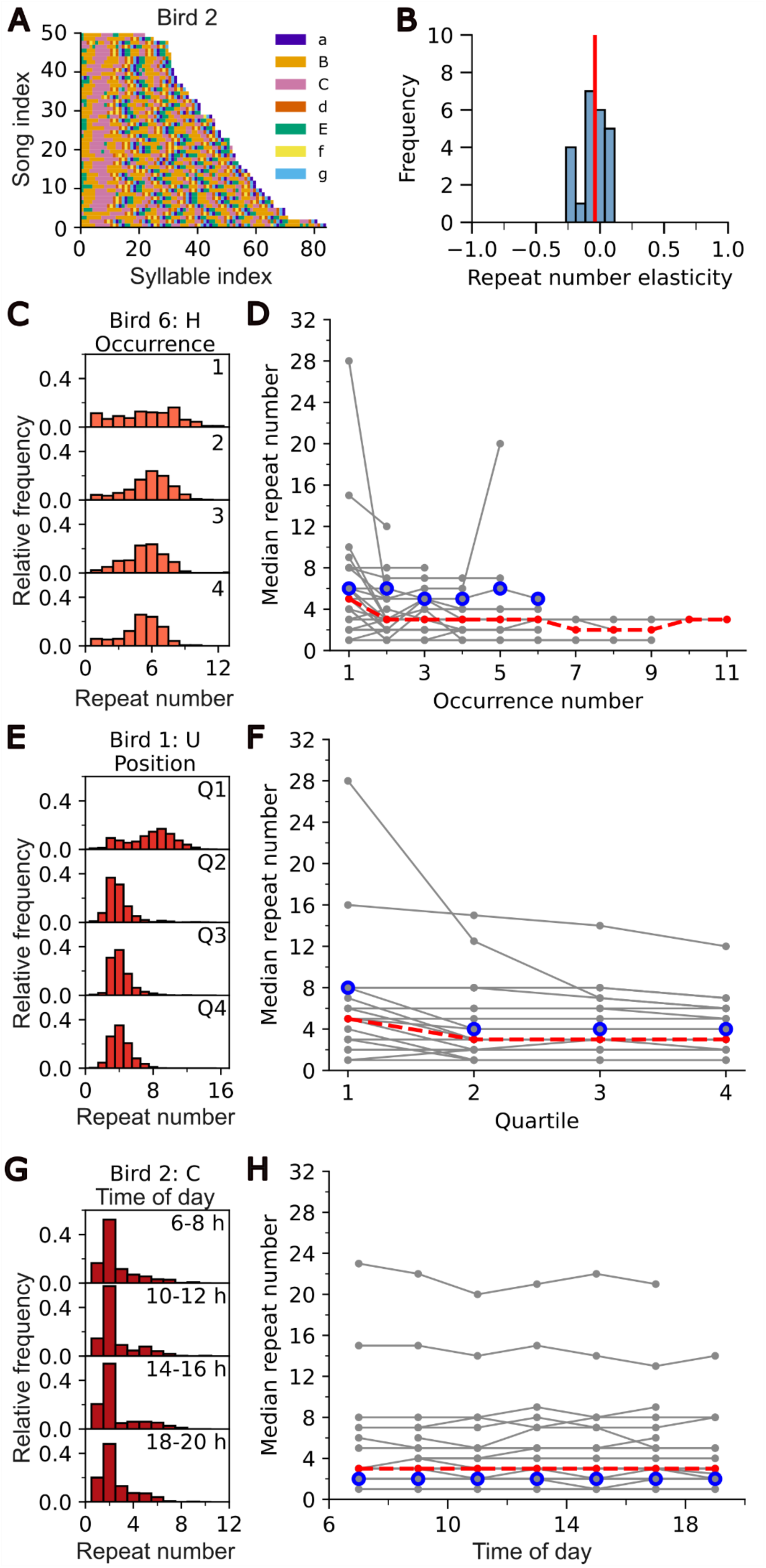
Repeat numbers are not strongly influenced by song bout length, occurrence number, position in song, and time of day. (A) 50 randomly sampled songs from bird 2 colour-coded by phrase type and sorted in increasing order of bout length. Uppercase and lowercase letters in the legend correspond to repeat phrases and non-repeating syllables respectively. (B) Repeat number elasticities (stretch proportional to bout length, see Methods) of all repeat phrases (n = 23 phrases from 6 birds). Red line indicates median. (C) Repeat distributions of example phrase H from bird 6 for first 1-4 occurrences in the bout. (D) Median repeat numbers of all phrases from 6 birds at different occurrence numbers. Points for the same phrase type are connected with grey lines; examples shown on the left are circled in blue. Red line indicates median across all phrases. (E) Repeat distributions of example phrase U from bird 1 in different quartiles of the bout. (F) Median repeat numbers of all repeat phrases from 6 birds in different quartiles of the bout. (G) Repeat distributions of example phrase C from bird 2 for four different two-hour time windows during the day. (H) Median repeat numbers of all phrases from 6 birds at different times of the day.

### Repeat numbers can be influenced by bout-level factors and time of day

Correlations between repeat numbers of phrases could be explained by other factors operating at the level of song bouts or the whole day. For example, if all phrases produced later in the bout or later in the day were longer or shorter due to factors like motivation, warm-up or fatigue, this could lead to correlations between pairs of phrases. We investigated this by analysing whether repeat numbers are influenced by the repeated occurrence of the same phrase in a bout (occurrence number), its relative position in the song bout, and time of day. All analysed phrases showed a significant effect of occurrence number (**Figures 3C, D**; p < 0.05 for 23/23 phrases), but this effect was largely idiosyncratic to each phrase, with slopes of the regression lines ranging from −1.45 to 0.10 with a mean of −0.25 ± 0.07. Similar to the effect of occurrence number, 22/23 phrases (95.6%) showed a significant effect of relative position in the bout, with the exact effect largely idiosyncratic to each phrase (**Figures 3E, F**; slopes of the regression lines ranged from −2.23 to 0.60 with a mean of −0.88 ± 0.19). Conversely, time of day had only small effects on repeat numbers, with a significant influence on 9/23 (39.1%) phrases analysed (**Figures 3G, H**) and slopes of the regression lines ranging from −0.08 to 0.04, mean 0.00 ± 0.01.

### Links between repeat phrases are not explained by syllable acoustics

Could factors like muscle fatigue or warm-up explain the correlation patterns? We reasoned that if syllables with similar acoustics rely on similar vocal production mechanisms, then the repeated production of one could impose constraints on the production of the other [43–45]. There was no significant difference between syllable acoustic distances across groups of positively, negatively or uncorrelated pairs of phrases, either when focusing on adjacent phrases or on all phrase pairs (**Figures S2 D,E**).

### Identity and repeat number of sequence context are the most important factors predicting repeat numbers

Since the identities and repeat numbers of surrounding syllables, as well as other factors such as bout length, position in the song bout and time of day could influence repeat numbers, we sought to weigh the relative importance of these factors using random forest models as illustrated in **Figure 4A**. We used SHAP (Shapley Additive Explanations) values to rank the relative importance of identity and repeat number of sequence context for up to four surrounding phrases, and of other factors in predicting the repeat number of each phrase. **Figure 4B** shows the normalised SHAP importances of an example model, which are broadly consistent with correlation results for the same example phrase (inset). Overall, we were able to fit good models to sixteen phrases from five birds (see Methods) and observed that the identities, as well as repeat numbers of preceding and following phrases were the most important factors in predicting repeat numbers, with trends towards decreasing importance as the distance from the modelled phrase increased. Other factors such as occurrence number, relative position, and time of day played an additional role for several phrases. Because the group effects of phrases at distances 3 and 4 were small, we only show two preceding and upcoming phrases for compactness (full model in **Figure S3**). Therefore, dependencies across phrases and the entire bout affect repeat numbers, suggesting that hierarchical motor organisation of Bengalese finch song may be necessary to explain these links.

**Figure 4:**
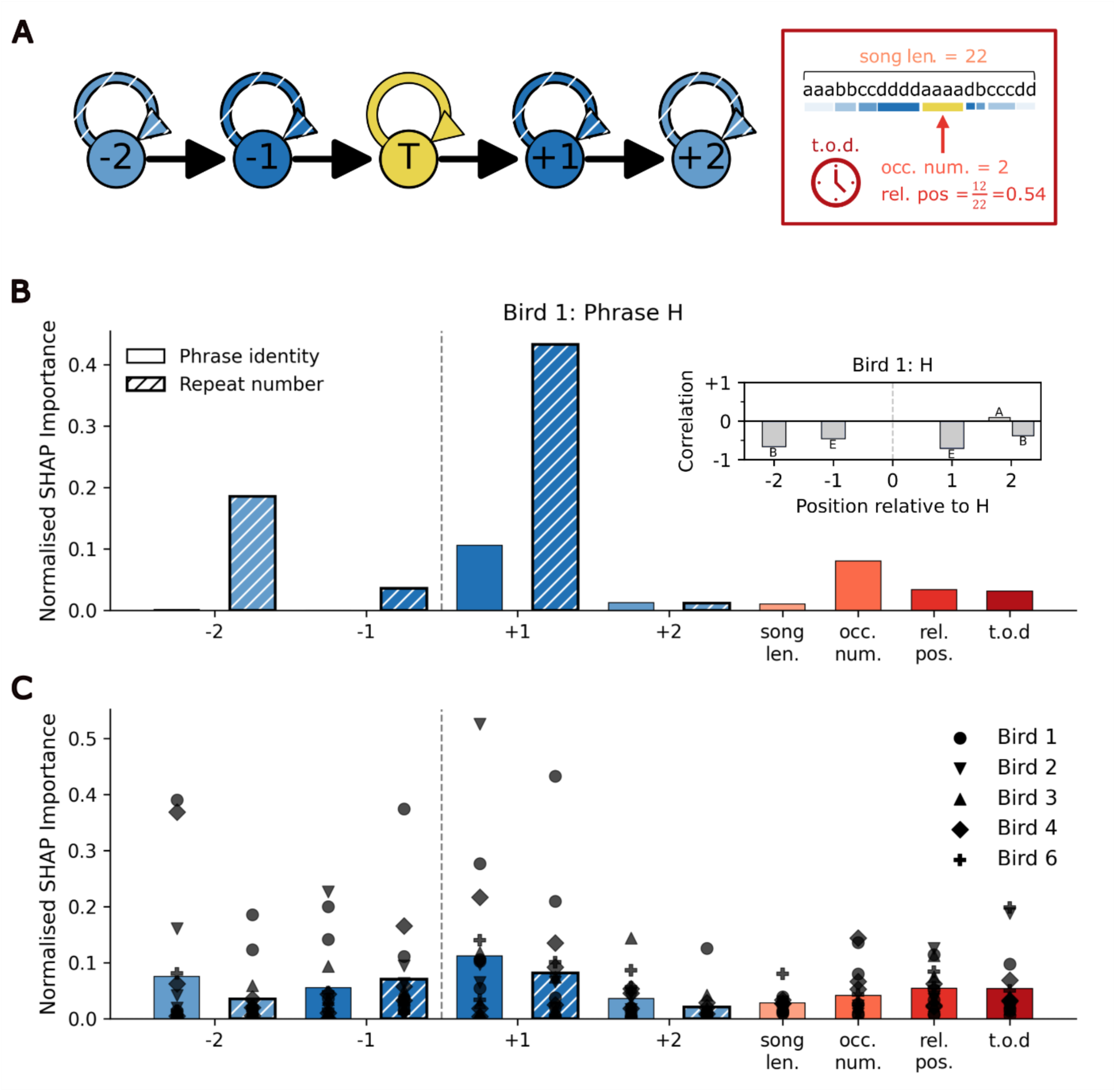
Random forest models indicate that repeat numbers are influenced by preceding and upcoming sequence contexts in the song. (A) Schematic representation of the random forest model. The model predicts the repeat number of example target phrase T (indicated in yellow) based on the identities (filled) and repeat numbers (hatched) of surrounding phrases. Contexts beyond two preceding and upcoming phrases are excluded for visualisation. The red box on the right shows other factors used as features in the model. Song len.: song bout length; t.o.d.: time of day; occ. num.: occurrence number; rel. pos.: relative position (B) Normalised SHAP importances from the random forest model for phrase H of bird 1. Hatched bars indicate SHAP importance for repeat number and blue unhatched bars indicate SHAP importances for identity of the sequence context. Other factors are depicted in red, abbreviations as in A. Inset shows correlation values for the same phrase, as in **Figure 2D**. (C) Normalised SHAP importances from random forest models for 16 phrases from 5 birds. Colours, abbreviations and shading as in panels A,B. Different phrases from the same birds are indicated by matching shapes.

## Discussion

In Markov chain models of song production, repeat phrases are modelled as self-transitions in the syllable sequence and rely on local stochastic decisions after each syllable to determine whether to continue the phrase or to move on to the next phrase [23–26,28]. Crucially, such a local, stochastic mechanism precludes non-local dependencies between distant phrases in the song bout, which we report here. Our results challenge a Markovian understanding of sequence generation in Bengalese finches and instead suggest a song organisation where the production of specific pairs of adjacent or distant phrases is linked [46,47].

How might links between distinct phrases, such as the correlated A-U phrases in **Figure 2** arise? Markov models of the syllable sequence correspond to a neuronal production mechanism in the well-studied song circuit [48–50] whereby networks of premotor neurons in nucleus HVC (proper name) control the production of individual syllables [51–54], and the next state is chosen through a stochastic winner-take-all mechanism that plays out at the end of each syllable [23,24,27,28,55]. Realistic repeat distributions for Bengalese finch song can be generated using a model with adapting self-transition probabilities, possibly due to adaptation of auditory feedback that sustains syllable repetition [28]. To capture correlations across phrases, such models would need to be extended by modelling distinct repeat phrases as reliant on a joint feedback parameter, so that singing one syllable may adapt the other.

Alternatively, repeat phrases may not be generated by a stochastic decision after every single syllable, but rather the approximate time spent in the phrase could be predetermined on any given rendition, with a different mechanism filling the allotted time with syllables. Such a mechanism has recently been reported for generating repeated vocalisations in the singing mouse [56], and would be consistent with firing patterns of neurons in song control nucleus HVC, which exhibit ramping bursts of activity encoding the relative progress through the repeat phrase [57]. The finding by Fujimoto et al. suggests that the approximate length of a repeat phrase is determined at the beginning of the phrase, setting the slope of the ramping bursts and generating longer phrases with shallow ramps on some renditions and short phrases with steep ramps on other renditions. Correlation patterns in this model could be explained by linked phrases sharing a joint ramping parameter and thus following steeper or shallower ramping patterns in the same bouts. In either production model, links between distant phrases suggest the presence of higher order structure in Bengalese finch song, with joint control of adaptation or ramping in distinct phrase types across the bout.

Important follow-up experiments should aim to test these links with behavioural interventions. For example, if different repeat phrases are controlled by the same auditory adaptation factor, then interrupting one phrase by disturbing auditory feedback [58] may likewise shorten other phrases linked with it. On the other hand, if higher structure in the motor production mechanism explains observed correlations, interrupting the execution of one phrase might leave its linked phrases undisturbed. Similarly, training birds to change repeat numbers through reinforcement learning [29,31,59] could test whether linked phrases can be manipulated separately, or whether learning to change the length of one will generalize to the length of the other [60,61].

What neuronal mechanisms could contribute to the observed links? If the lengths of repeat phrases are determined by stochastic premotor networks in HVC, links in the control of distinct phrases could arise developmentally, for example through incomplete splitting of syllable networks from precursor networks during the song learning period [62–64]. This would lead to retained links in the neuronal control of acoustically distinct adult syllables. Characterisation of the premotor activity underlying song production in HVC [52,63] in Bengalese finches could reveal whether correlated syllables share more similar premotor networks than unrelated syllables [65]. Alternatively, the anterior forebrain pathway, a basal-ganglia pathway involved in song learning, has been suggested to contribute to the initiation, termination and control of repeating syllables [32,57,66–68]. For example, lesions or inactivation of Area X, a songbird basal ganglia nucleus, can lead to an increase in syllable repetition in both Bengalese finches and zebra finches [66–68]. Similarly, lesions of mMAN (medial magnocellular nucleus of the anterior nidopallium), an output of the medial part of the anterior forebrain pathway, make repeat numbers of repeat phrases more variable [32] and pharmacological manipulations of lMAN (lateral magnocellular nucleus of the anterior nidopallium) the output of the lateral anterior forebrain pathway, increase syllable repetition in Bengalese finches [67]. Finally, the influence of other factors such as time of day or position in the bout could hint at additional mechanisms downstream of the forebrain song control system, such as fatigue [43,69] or respiratory constraints [45] in determining the lengths of repeat phrases.

This study adds to a growing body of evidence for long-range dependencies in Bengalese finch songs [16,57,70–72]. Similarly, in canary song, transition probabilities, the lengths of repeat phrases and premotor HVC activity can depend on the preceding contexts [41,73]. At a higher hierarchical level, songbird species with multiple song types can exhibit even longer-range temporal dependencies in the sequential organisation of different song types [74,75]. Elucidating the neuronal mechanisms underlying linked repetition will provide important insights into the production of such hierarchical and complex vocal sequences.

## Code Accessibility

All code necessary for analysing data and creating figures related to this manuscript can be accessed at https://github.com/pbinwal/BV_2026.

## Acknowledgements

This work was supported by the Deutsche Forschungsgemeinschaft (DFG, German Research Foundation) – Project number 536953998 and 532521431. We thank Avani Koparkar for help with data collection and code for acoustic analyses, and members of the Veit lab Jacqueline Göbl, Lioba Fortkord, Franziska Heubach and Abhilipsa Das for comments on the manuscript. We acknowledge the use of AI technologies (Claude and Chatgpt) for assistance with coding.

## Conflict of interest statement

The authors declare no competing financial interests.

## Supplementary Materials

**Figure S1.**
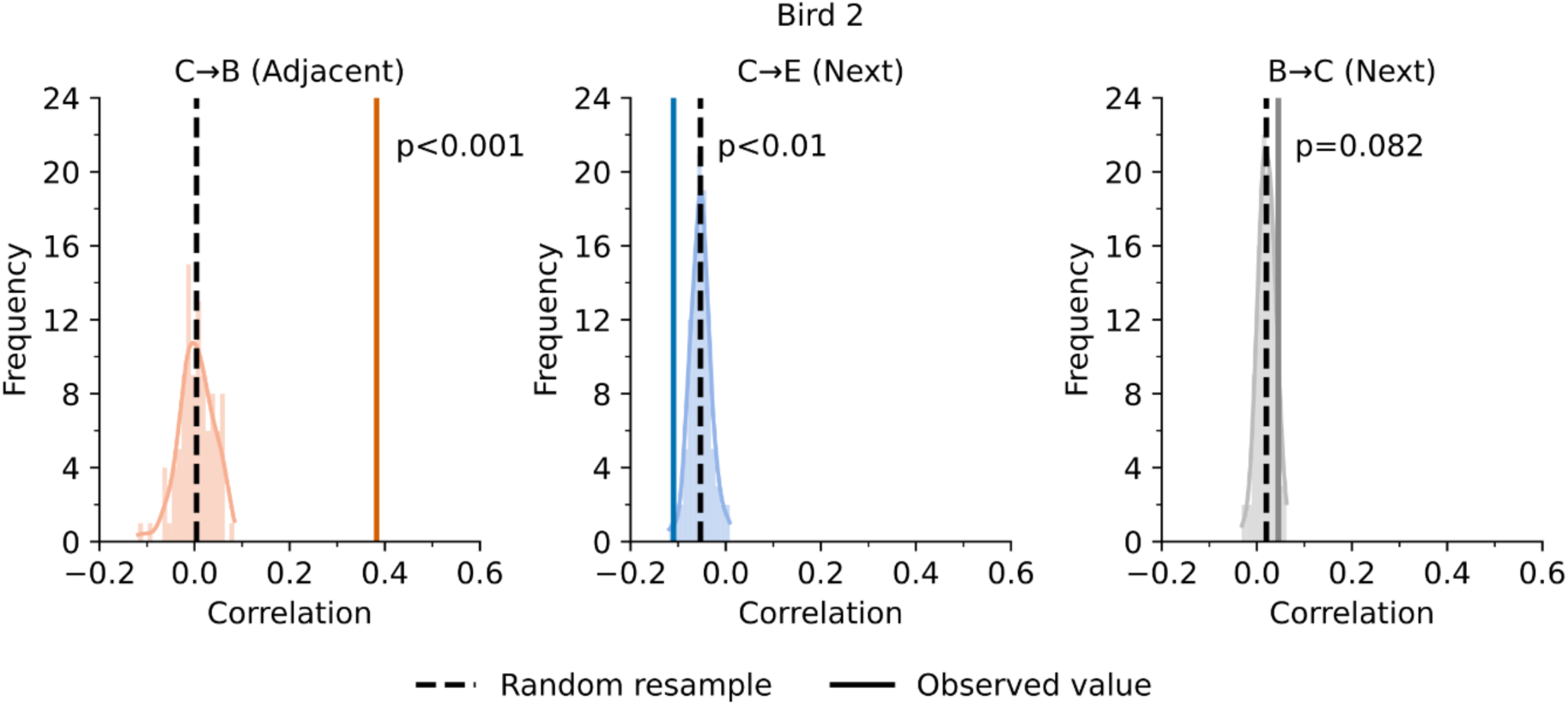
Correlations between repeat phrases are not observed in resampled data and are therefore not explained by constraints imposed by repeat distributions of phrases. (A) Example distributions of correlation values in resampled data from bird 2 also shown in **Figure 2D-F**; p-values from one-tailed z-tests.

**Figure S2.**
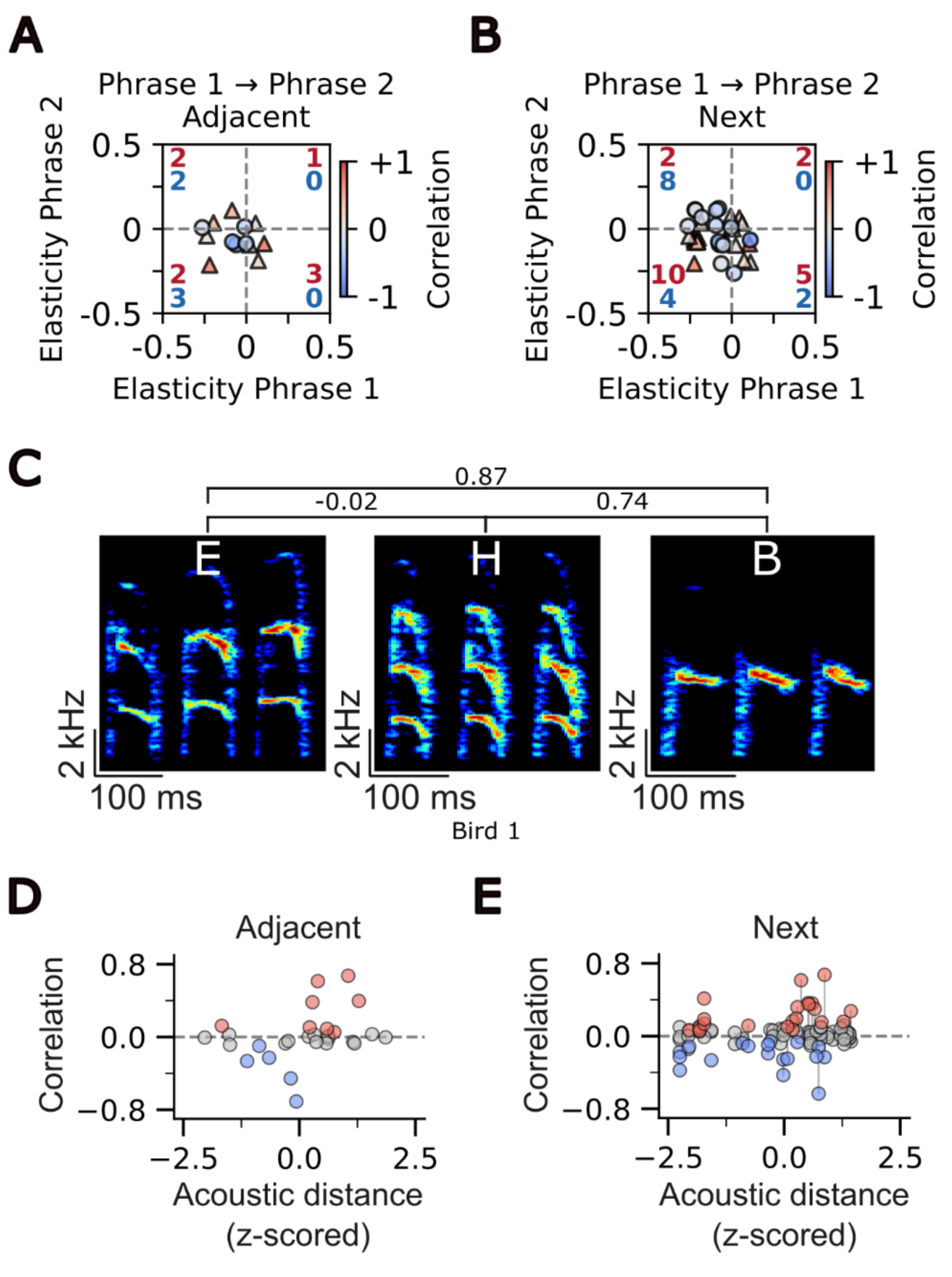
Repeat number elasticities and syllable acoustics do not explain observed correlations. (A) Elasticities of adjacent correlated phrase pairs. Triangles and circles represent positively and negatively correlated pairs, respectively. Red and blue numbers represent the counts of positively and negatively correlated pairs in each quadrant. 37.5% and 62.5% of positively correlated adjacent phrase pairs had the same or opposite elasticity signs, respectively. 60% and 40% of negatively correlated adjacent pairs had the same or opposite elasticity signs, respectively. (B) Elasticities of all correlated phrase pairs. Of all positively correlated pairs, 63.2% and 36.8% had the same or opposite elasticity signs, respectively. For negatively correlated adjacent pairs, 28.6% and 71.4% had the same or opposite signs, respectively. Shapes, colours and numbers as in panel A. (C) Acoustic distances (z-scored) between three example syllables from Bird 1. (D) Scatter plot depicts correlations and acoustic distances between adjacent phrase pairs. Positive, negative, and non-significant correlations are depicted as red, blue, and grey circles respectively. (E) Scatter plot depicts correlations and acoustic distances between all phrase pairs. Colours as in panel D. Vertical grey lines connect reverse pairs (for example A→B and B→A) which, by definition, had the same acoustic distance when considering all occurrences (see Methods).

**Figure S3.**
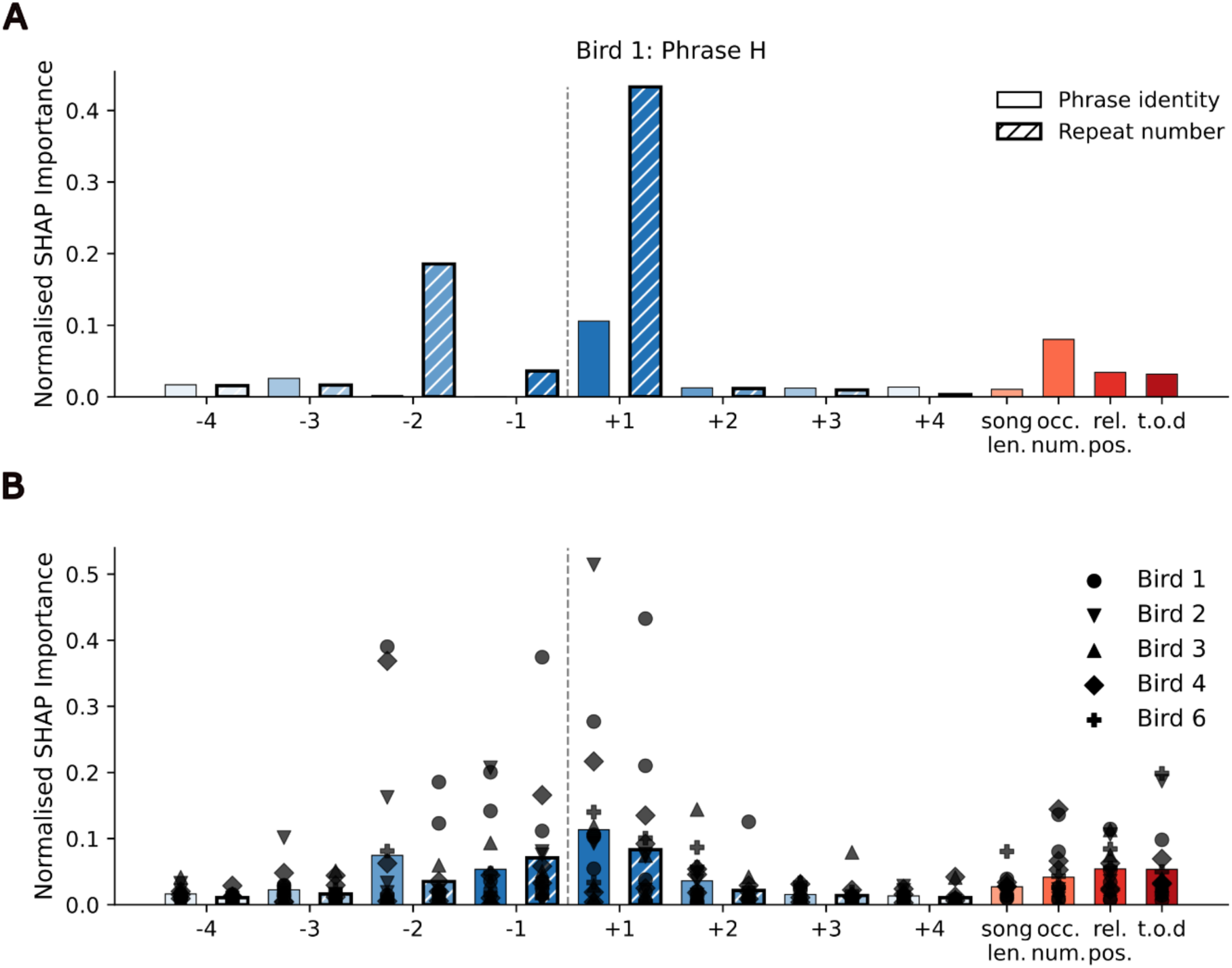
Full data including all sequence contexts for the models shown in Figures 4B, C. Full sequence context plots for the example model and all models shown in **Figures 4B, C**. (A) Bar plot depicts the normalised SHAP importances from the random forest model for phrase H of bird 1 including sequence contexts from −4 to +4. Hatched bars with thicker outlines and unhatched bars in blue indicate the SHAP importances for the sequence context and repeat numbers of the context, respectively. Other factors are depicted in shades of red. (C) Bar plot depicts the normalised SHAP importances from random forest models for 16 phrases from 5 birds. SHAP importances from phrases belonging to different birds are depicted by different shapes. Colours and bar shading as in panel A.

## Notes

### Competing Interest Statement

The authors have declared no competing interest.

